# Hippocampal Interneuronal Dysfunction and Hyperexcitability in a Porcine Model of Concussion

**DOI:** 10.1101/2022.03.08.483543

**Authors:** Alexandra V. Ulyanova, Christopher D. Adam, Carlo Cottone, Nikhil Maheshwari, Michael R. Grovola, Oceane E. Fruchet, Jami Alamar, Paul F. Koch, Victoria E. Johnson, D. Kacy Cullen, John A. Wolf

## Abstract

Cognitive impairment is a common symptom following mild traumatic brain injury (mTBI or concussion) and can persist for years in some individuals. Hippocampal slice preparations following closed-head, rotational acceleration injury in swine have previously demonstrated reduced axonal function and hippocampal circuitry disruption. However, electrophysiological changes in hippocampal neurons and their subtypes in a large animal mTBI model have not been examined. Using *in vivo* electrophysiology techniques, we examined laminar oscillatory field potentials and single unit activity in the hippocampal network 7 days post-injury in anesthetized minipigs. Concussion altered the electrophysiological properties of pyramidal cells and interneurons differently in area CA1. While the firing rate, spike width and amplitude of CA1 interneurons were significantly decreased post-mTBI, these parameters were unchanged in CA1 pyramidal neurons. In addition, CA1 pyramidal neurons in TBI animals were less entrained to hippocampal gamma (40 - 80 Hz) oscillations. Stimulation of the Schaffer collaterals also revealed hyperexcitability across the CA1 lamina post-mTBI. Computational simulations suggest that reported changes in interneuronal physiology may be due to alterations in voltage-gated sodium channels. These data demonstrate that a single concussion can lead to significant neuronal and circuit level changes in the hippocampus, which may contribute to cognitive dysfunction following mTBI.

## Introduction

Concussion, or mild TBI (mTBI), is a prevalent form of brain injury, affecting approximately 1.3 million people every year in the US alone (Bazarian et al., 2005). Cognitive impairment is common following concussion and deficits in episodic memory, learning, and delayed recall have been described (Anita et al., 2008; de Freitas Cardoso et al., 2019; Giza et al., 2018). Importantly, a growing body of data suggests that neurocognitive dysfunction observed post-concussion can persist for years after initial insult in some people (King and Kirwilliam, 2011; Levin et al., 2021; Rabinowitz et al., 2015; Røe et al., 2009; Sterr et al., 2006). Given the important role of the hippocampus in learning and memory, it is a central region of interest in diseases affecting cognitive processes (for review, see (Carr et al., 2011)). While the underlying mechanisms of cognitive dysfunction following mTBI remain largely unknown, structural changes in the human hippocampus up to 1-year post-concussion have been reported using imaging techniques (Singh et al., 2014; Strain et al., 2015). Recent reports also link concussion to hyperexcitability (King et al., 2019; Major et al., 2015). Moreover, recent reports based on the epidemiological data may indicate a 2-times increased risk for post-traumatic epilepsy in adult population with mTBI (Annegers and Coan, 2000; Lolk et al., 2021; Pugh et al., 2015).

Rodent studies of TBI have shown changes in neuronal firing and local hippocampal oscillations, potentially suggesting abnormalities in temporal coding, a crucial element of episodic memory formation and encoding (Broussard et al., 2020; Ondek et al., 2020; Paterno et al., 2015). While studies utilizing rodent models of TBI can provide some mechanistic insight into electrophysiological changes following TBI, brains of large animals such as pigs, with their gyrencephalic structure and appropriate white-to-grey matter ratios, more closely resemble human architecture and are important for an accurate biomechanical modeling of all aspects of human TBI (for review, see (Johnson et al., 2013)). Using a biomechanically relevant and gyrencephalic model of concussion in swine, we demonstrated injury-related compensatory mechanisms and a reduction in axonal function in the *ex vivo* hippocampus following TBI in the apparent absence of intra-hippocampal neuronal or axonal degeneration (Wolf et al., 2017). Deafferentation of temporal lobe structures (such as the hippocampus) via axonal pathology has been proposed as a driver of hyperexcitability in this model and may disrupt the precise timing of hippocampal oscillations and associated neuronal activity (for review, see (Wolf and Koch, 2016)). Diffuse axonal injury is the predominant pathology observed in this preclinical model, however direct evidence that concussion disrupts neuronal network activity, specific neuronal subtypes, or laminar oscillations in the hippocampus of the intact swine has yet to be demonstrated.

We utilized an established porcine model of closed-head, rotational acceleration to induce concussion, and then applied high-density *in vivo* electrophysiology techniques to examine neuronal and oscillatory changes in the intact hippocampus at 7 days post-injury. Notably, the function of hippocampal circuits was assessed using laminar, multichannel silicon probes specifically designed for recording from deep brain structures of large animals (Ulyanova et al., 2019; Ulyanova et al., 2018). Laminar oscillatory field potentials and single unit activity were examined and compared between injured and control animals under isoflurane anesthesia. Our data suggest that even a single concussion may alter both individual neuronal function and circuit-level activity in the hippocampus and may provide insight into aspects of hyperexcitability and cognitive dysfunction following concussion.

## Results

### Concussion Affects Electrophysiological Properties of Hippocampal CA1 Neurons

To examine changes in hippocampal circuitry induced by concussion, *in vivo* electrophysiological recordings were obtained using 32-channel silicon probes that spanned the laminar structure of CA1 in the dorsal hippocampus (Figure 1A). These probes allowed recording of local field potential across layers of CA1 and isolation of single units using available spike sorting algorithms (see Methods) (Gold et al., 2006). Electrophysiological properties of CA1 neurons such as firing rate, spike width and amplitude (Figure 1B) were calculated and compared between control (n_animals_ = 8; n_cells_ = 125) and post-mTBI (n_animals_ = 9; n_cells_ = 93) animals at 7 days post-concussion (Figure 1C). While the number of recorded cells per animal was not different between the groups (control = 16 ± 3 cells vs. post-mTBI = 10 ± 3 cells, p = 0.2397), the firing rate, spike width, and spike amplitude were all significantly reduced in post-mTBI animals (Figure 1C). The firing rate of 4.43 ± 0.50 Hz in the control animals was reduced to 2.09 ± 0.27 Hz in the post-mTBI animals (p = 0.0002). The spike width of 0.358 ± 0.015 msec in the control group was significantly reduced to 0.270 ± 0.008 msec in the post-mTBI group (p < 0.0001), and the spike amplitude was significantly reduced from 300 ± 27 µV in the control group to 187 ± 27 µV in the post-mTBI group of animals (p = 0.0003). These findings suggest that concussion may alter the intrinsic properties of CA1 neurons, which could affect their synchronization with hippocampal oscillations.

**Figure 1.**
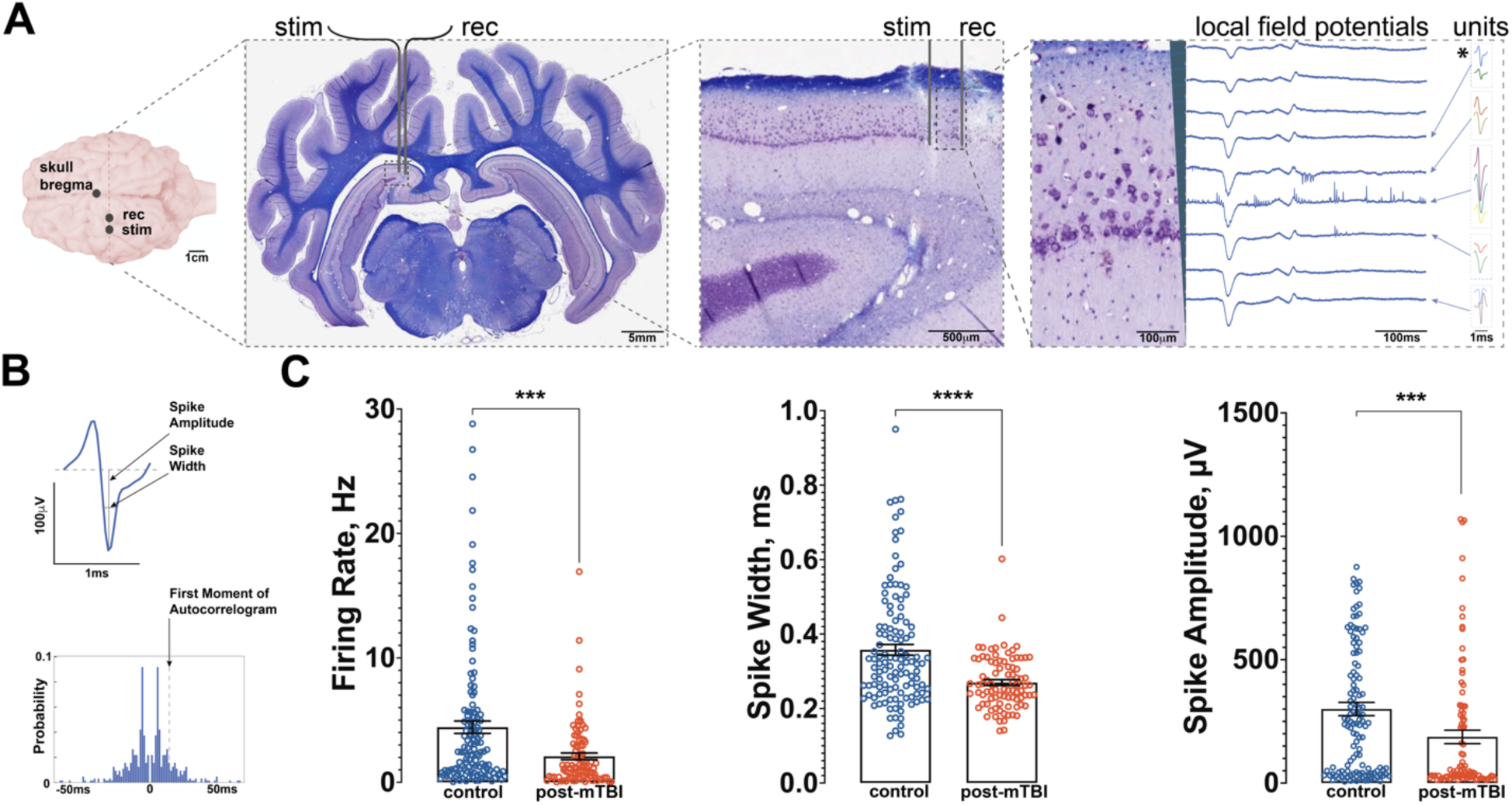
Concussion Affects Electrophysiological Properties of Hippocampal CA1 Neurons. **A)** Anatomical locations for recording and stimulation electrodes depicted relative to bregma point (scale bar = 1cm). A representative coronal section through the plane of recording/stimulation showing hippocampal anatomy, stained with LFB/CV as previously described (Ulyanova et al., 2018) (scale bar = 5mm). Stimulation electrode and recording electrode shown overlaid on dorsal hippocampus (scale bar = 500μm). Local field potentials (scale bar = 100ms) and multiple single units (scale bar = 1ms) recorded with 32-channel laminar probe (scale bar = 100μm). Note multiple single units present on the same channel (boxed and color coded). **B)** A representative single unit (blue, depicted with * in A), displaying electrophysiological parameters such as firing rate, spike width, and spike amplitude (top). First moment of autocorrelogram was also calculated for each spike (bottom). **C)** At 7 days post-mTBI, the firing rate of hippocampal CA1 neurons was significantly reduced in the control (4.43 ± 0.50 Hz) vs. post-mTBI (2.09 ± 0.27 Hz) groups of animals (p = 0.0002). In addition, the spike width of 0.358 ± 0.015 msec recorded in the control animals was significantly reduced to 0.270 ± 0.008 msec in the post-mTBI animals (p < 0.0001). The spike amplitude was also significantly reduced from 300 ± 27 µV in the control animals to 187 ± 27 µV in the post-mTBI animals (p = 0.0003). Data presented as mean ± SEM.

### Pyramidal Neurons in CA1 Preserve Their Firing Properties Following Concussion

To further characterize the effects of mTBI on the hippocampal circuitry and to assess changes in specific subpopulations of cells in CA1, recorded cells were separated into putative CA1 pyramidal cells or putative CA1 interneurons (Wheeler et al., 2015). Previously published criteria for rodent hippocampal cells were used for classification (Broussard et al., 2020; Csicsvari et al., 1999), with several modifications accounting for species differences. Briefly, the probe’s anatomical location coupled with electrophysiological characteristics were utilized to exclude cells recorded at the bottom portion of the probe, as these cells could be identified as putative dentate granule cells, mossy cells, or interneurons in the dentate (control: n_cells_ = 14, post-mTBI: n_cells_ = 5). Additionally, unclassifiable cells located throughout the length of the probe (control: n_cells_ = 10, post-mTBI: n_cells_ = 16) were omitted from the analysis. Next, electrophysiological parameters (firing rate, spike width, and first moment of autocorrelogram, Figure 1B, 2A, 3A) of the remaining single units were used to classify the remaining neurons (see Methods for more details) as either putative CA1 pyramidal cells (control: n_pyr_ = 49, post-mTBI: n_pyr_ = 37; Figure 2) or putative CA1 interneurons (control: n_int_ = 52, post-mTBI: n_int_ = 38; Figure 3). The number of either pyramidal cells (control = 7 ± 2 cells vs. post-mTBI = 4 ± 1 cells) or interneurons (control = 6 ± 2 cells vs. post-mTBI = 4 ± 2 cells) per animal was not different between the control and post-mTBI groups (CA1 pyramidal cells: p = 0.3515; CA1 interneurons: p = 0.4314).

**Figure 2.**
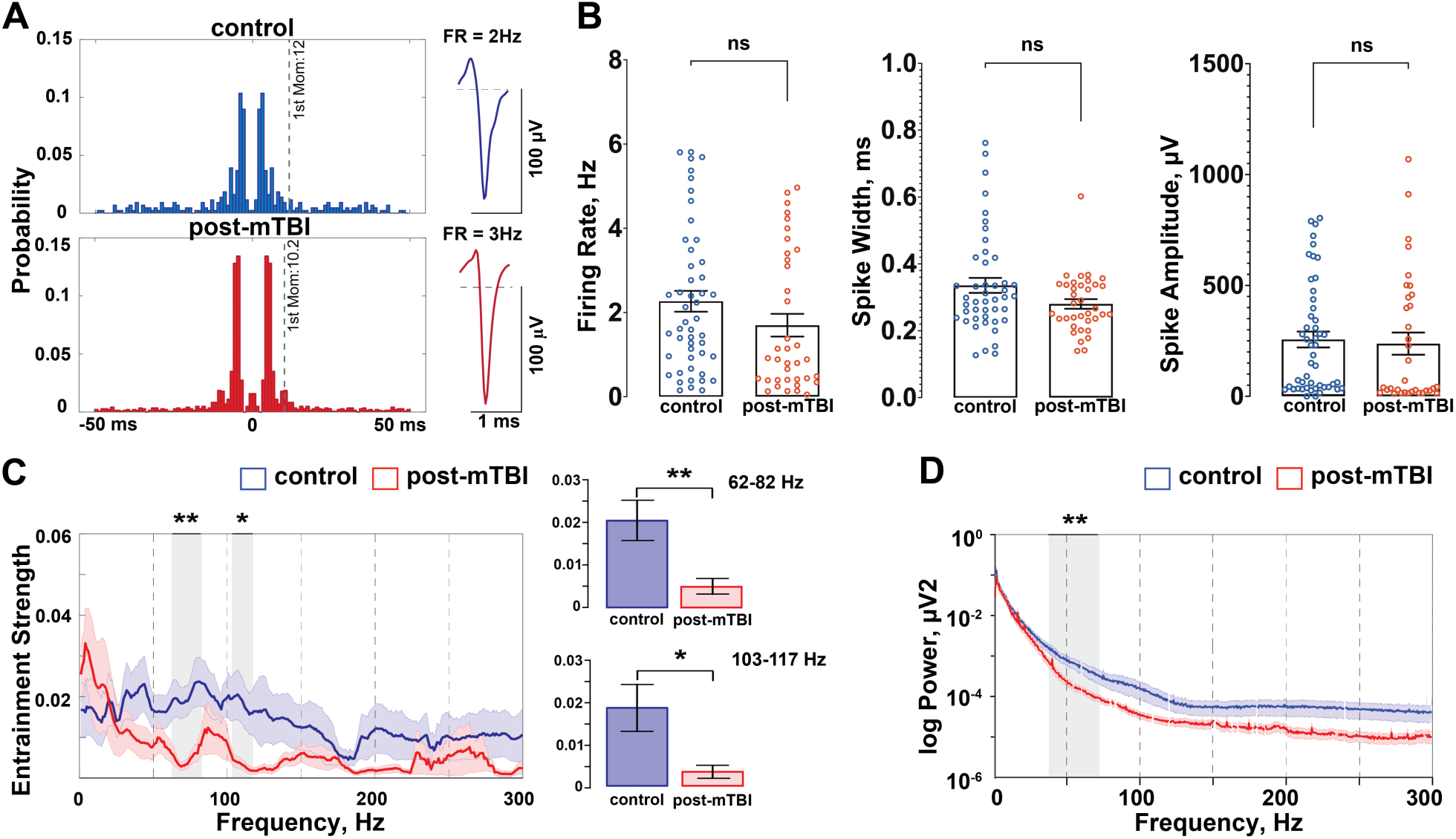
Pyramidal Neurons in CA1 Preserve Their Firing Properties Following Concussion but Are Less Entrained to Gamma. **A)** Representative examples of putative CA1 pyramidal cells from the control (blue) and post-mTBI (red) animals are shown. **B)** Electrophysiological properties of pyramidal cells were not changed 7 days following mTBI. The parameters analyzed include firing rate (control = 2.27 ± 0.25 Hz, vs. post-mTBI = 1.70 ± 0.27 Hz; p = 0.0555), spike width (control = 0.336 ± 0.022 msec vs. post-mTBI = 0.280 ± 0.014 msec; p = 0.2269), and spike amplitude (control = 256 ± 36 µV vs. post-mTBI = 238 ± 50 µV; p = 0.1372). Data presented as mean ± SEM. **C)** Entrainment of CA1 pyramidal cells to local hippocampal oscillations was significantly decreased in the 62 - 82 Hz (p < 0.01) and the 103 - 117 Hz (p < 0.1) frequency ranges in the post-mTBI animals. **D)** Power spectral density analysis shows a significant decrease in the 35 – 75 Hz frequency range (p < 0.01) following mTBI. Data presented as mean ± STD. Noise associated with 60-Hz cycle was removed manually.

**Figure 3.**
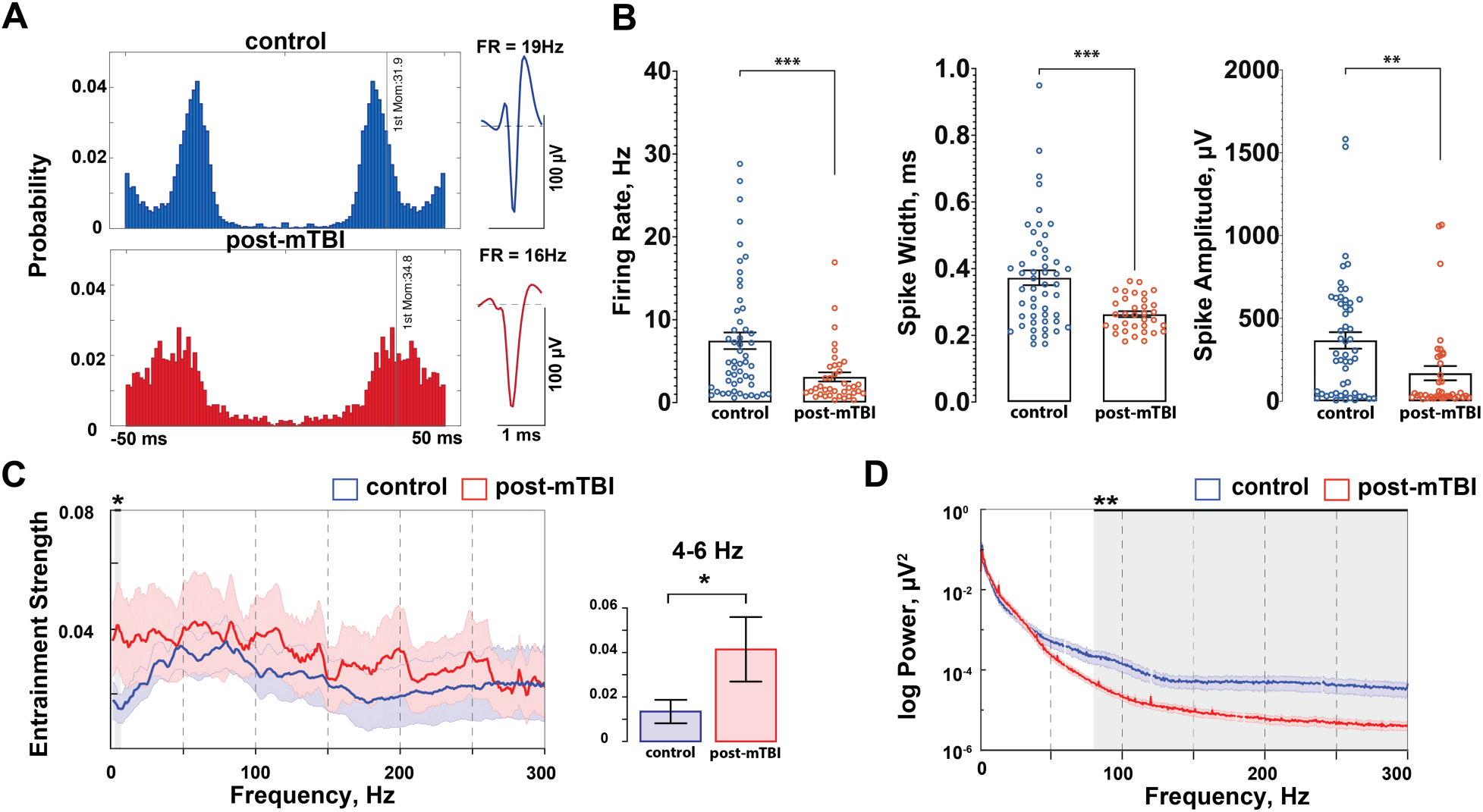
Interneurons in CA1 Are Preferentially Affected Following Concussion. **A)** Representative examples of CA1 interneurons from control (blue) and post-mTBI (red) animals. **B)** Electrophysiological properties of CA1 interneurons were significantly altered 7 days post inertial injury. Firing rate of CA1 interneurons was significantly reduced from 7.46 ± 1.0 Hz in control animals (blue) to 3.09 ± 0.55 Hz (red) in post-mTBI animals (p = 0.0008). The spike width was significantly reduced from 0.373 ± 0.022 msec in control animals (blue) to 0.263 ± 0.009 msec in post-mTBI animals (red) (p = 0.0004). Spike amplitude was also significantly reduced from 367 ± 50 µV in control animals (blue) to 170 ± 43 µV in post-mTBI animals (p = 0.0036). Data presented as mean ± SEM. **C)** Entrainment of CA1 interneurons to local hippocampal oscillations was significantly increased in the 4 - 6 Hz frequency range (p < 0.05). **D)** Power of high frequency oscillations was significantly decreased in the 80 - 300 Hz frequency range (p < 0.01). Data presented as mean ± STD. Noise associated with 60-Hz cycle was removed manually.

While some changes in the electrophysiological properties of putative CA1 pyramidal cells in the post-mTBI group trended towards significance (e.g., firing rate or spike width), firing rate (control = 2.27 ± 0.25 Hz vs. post-mTBI = 1.70 ± 0.27 Hz; p = 0.0555), spike width (control = 0.336 ± 0.022 msec vs. post-mTBI = 0.280 ± 0.014 msec; p = 0.2269), and spike amplitude (control = 256 ± 36 µV vs. post-mTBI = 238 ± 50 µV; p = 0.1372) were not significantly different between the control (n = 8) and post-mTBI (n = 9) groups of animals at 7 days following concussion (Figure 2B). Recent studies in rodent models of TBI have shown changes in both neuronal firing and hippocampal local field potentials, suggesting abnormalities in temporal coding, a process through which tightly coordinated network activity seen in the local field potential controls the precise timing of neuronal firing (Broussard et al., 2020; Ondek et al., 2020; Paterno et al., 2015). Therefore, to examine whether CA1 pyramidal cells continue to interact with hippocampal oscillations properly following concussion, entrainment of CA1 pyramidal cells to local field potentials was also investigated (Figure 2C). Notably, entrainment strength (a measure that determines how tightly coupled neuronal firing is to particular phases of an oscillation) of CA1 pyramidal cells to local oscillations was significantly reduced in the 62 - 82 Hz frequency band (control = 0.020 ± 0.005 vs. post-mTBI = 0.012 ± 0.005, p < 0.01) and in the 103 - 117 Hz frequency band (control = 0. 018 ± 0.006 vs. post-mTBI = 0.003 ± 0.001, p < 0.05). Additionally, we found a significant reduction in gamma oscillation (35 – 75 Hz) power (a measure that determines the strength of a particular oscillation in the overall EEG signal) detected locally to pyramidal CA1 cells (Figure 2D, p < 0.01). These results indicate that concussion disrupts the generation of gamma oscillations in the pyramidal cell layer due to the changes in afferent inputs coming into CA1 and/or alterations in local CA1 pyramidal-interneuron or interneuron-interneuron interactions.

### Inhibitory CA1 Interneurons Are Preferentially Affected Following Concussion

Analysis of putative CA1 interneurons revealed that their electrophysiological properties were significantly altered. In contrast to CA1 pyramidal cells, CA1 interneurons had a significantly reduced firing rate (control = 7.46 ± 1.0 Hz vs. post-mTBI = 3.09 ± 0.55 Hz, p = 0.0008), spike width (control = 0.373 ± 0.022 msec vs. post-mTBI = 0.263 ± 0.009 msec, p = 0.0004), and spike amplitude (control = 367 ± 50 µV vs. post-mTBI = 170 ± 43 µV, p = 0.0036) at 7 days post-concussion (Figure 3B). Interestingly, the firing rate of CA1 interneurons decreased as rotational velocity increased, changing significantly from 7.46 ± 1.0 Hz in control animals (n = 8) to 3.78 ± 0.93 Hz in animals injured at ∼190 rad/sec (188 ± 11 rad/sec, n = 4), and further to 2.33 ± 0.51 Hz in animals injured at ∼260 rad/sec (256 ± 7 rad/sec, n = 5) (p = 0.0030). Since the electrophysiological properties of CA1 pyramidal cells were not substantially altered (Figure 2B), these results indicate that CA1 interneurons are preferentially affected at 7 days following a single inertial injury. Notably, entrainment of CA1 interneurons to oscillations in the 4 - 6 Hz frequency range was significantly increased 7 days post-concussion (control = 0.013 ± 0.005 vs. post-mTBI = 0.041 ± 0.014, p < 0.05, Figure 3C). Additionally, there was a significant reduction in the power of high frequency oscillations (frequency range = 80 – 300 Hz) detected locally to interneurons (but not pyramidal cells) in CA1 cells, potentially due to the observed reduction in the firing rate of CA1 interneurons or desynchronization of the CA1 network (Figure 3D, p < 0.1). These results indicate that concussion alters intrinsic and/or synaptic properties of CA1 interneurons, which affects local gamma and high-frequency oscillations presumably generated by local CA1 interneurons. These changes likely alter the way CA1 pyramidal cells interact with local hippocampal oscillations.

### Hippocampal Hyperexcitability Post-Concussion

Reduced excitability of GABAergic inhibitory interneurons without a corresponding change in the activity of excitatory neurons in the hippocampus has been previously linked to hippocampal hyperexcitability (Hunt et al., 2011; Pavlov et al., 2011; Toth et al., 1997; Witgen et al., 2005). Moreover, the differential response to stimulation *ex vivo* has been previously reported between control and post-mTBI animals using this model (Wolf et al., 2017), further suggesting hyperexcitability in hippocampus post-concussion. Here, we used electrical stimulation of the Schaffer collateral inputs coming into the CA1 layer from CA3 in a subset of animals (control = 3, post-mTBI = 7) to assess changes in the excitability levels in the intact brain (also see (Ulyanova et al., 2018) for more details). Representative examples of the evoked laminar field potentials in dorsal CA1 using Schaffer collateral stimulation are shown as averaged responses over a 300 ms time period (20 stimulations, 2 s apart) for a control (blue, left panel) and a post-mTBI (red, right panel) animal (Figure 4A). During a series of the paired-pulse (PP) stimulations in the Shaffer collaterals (inter-pulse interval of 30 msec, 20 repeats, 5 sec apart), a sustained depolarization over 1 sec long, represented by negative deflection of the extracellular local field potentials (for review, see (Buzsáki et al., 2012)), was observed in a subset (n = 2 out of 5) animals injured at ∼260 rad/sec (Figure 4B, Box 1). Notably, this depolarizing shift (Figure 4C) was detected across all hippocampal layers and has features similar to the paroxysmal depolarizing shift previously observed in epileptic brain (for review, see (Hotka and Kubista, 2019)). In the example shown in Figure 4C, the depolarizing shift happened approximately 4 sec after the third PP stimulation and continued for over 1 sec, overlapping with the fourth PP stimulation. In addition, depolarization was followed by a loss of synaptic responses to PP stimulation (Figure 4B, Box 2). Other forms of hippocampal hyperexcitability such as paroxysmal rhythmic spikes were observed in the form of synchronized activity in the 6 - 8 Hz frequency range across all laminar layers in injured but not control animals (Figure 4D). Interestingly, spontaneous paroxysmal rhythmic spikes were also present during baseline recordings (no stimulation) in all post-mTBI (n = 9) but not control (n = 8) animals (not shown), potentially suggesting increased hyperexcitability and synchronization of hippocampal circuitry prior to the introduction of a stimulus.

**Figure 4.**
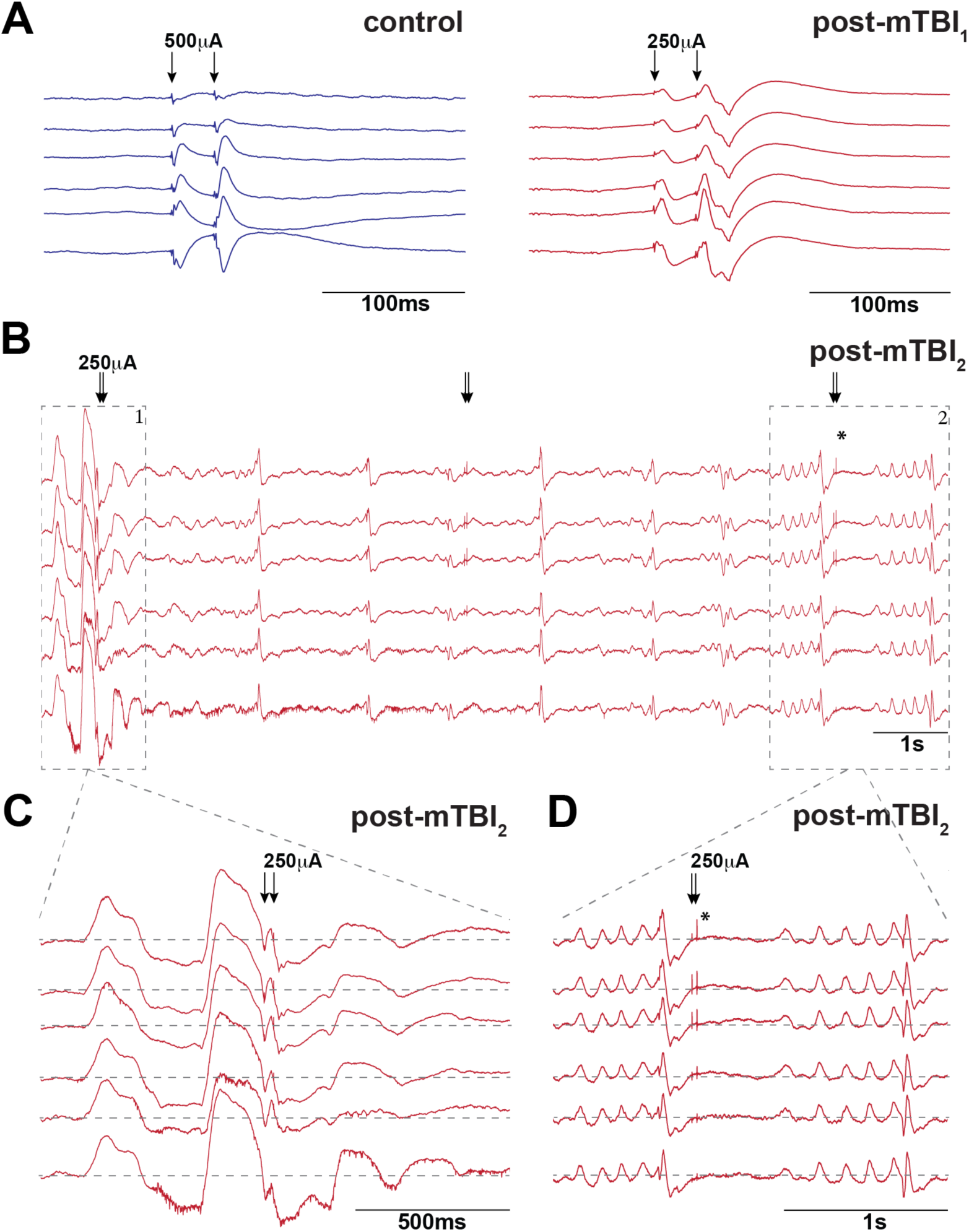
Hippocampal Hyperexcitability Following Concussion. **A)** Averaged CA1 laminar field responses to a train of paired pulse (PP) stimulation (inter-pulse interval, IPI = 50ms, 2s in between stimulations, 20 stimulations total) in Schaffer Collaterals (arrows) are shown for a control (stimulation amplitude = 500μA, left panel) and post-mTBI (stimulation amplitude = 250μA, right panel) animals. **B)** A depolarizing shift was observed in a subset (n = 2) of post-mTBI (but not control) animals during PP stimulation (IPI = 30ms, stimulation amplitude = 250μA, 5s in between stimulations, 20 stimulations total). In this example, the depolarizing shift happened approximately 4s following an application of 3^rd^ PP stimulation and continued for over 1s (Box1), overlapping with the 4^th^ PP stimulation (arrows). A prolonged depolarization followed by a loss of synaptic responses to PP stimulation (arrows) was also observed in all layers of hippocampus (Box 2, asterisk). **C)** Magnified view of Box 1 in B shows a period of prolonged depolarization as negative deflection over baseline, with multi-unit activity present on some channels (bottom two). **D)** Magnified view of Box 2 in B shows other forms of hippocampal hyperexcitability such as a loss of synaptic responses stimulation (asterisk) and paroxysmal rhythmic spikes in the form of synchronized activity in the 6-8 Hz frequency range across all laminar layers.

### Model of Parvalbumin-Positive (PV^+^) CA1 Interneurons Reveals Potential Dysfunction of Sodium Channels

Interneurons in the hippocampus are involved in the generation of gamma oscillations and provide input-specific inhibition to pyramidal cells via feedforward and feedback inhibition (Buzsáki and Wang, 2012; Mann et al., 2005; Soltesz and Deschenes, 1993). Specifically, it has been previously shown that parvalbumin-positive (PV^+^) basket cells provide peri-somatic, feedforward inhibition in the hippocampal CA1 region (for review, see (Klausberger and Somogyi, 2008; Pelkey et al., 2017)). These cells stabilize hippocampal circuitry during changes in input and control the selection of cell ensembles by allowing only certain number of pyramidal cells to fire in response to the stimulus (for review, see (Buzsáki and Wang, 2012; de Almeida Licurgo, 2009; Rennó-Costa et al., 2019)). Interestingly, pathological changes in PV^+^ interneurons in CA1 have been also shown to play a crucial role in the development of epilepsy (Ogiwara et al., 2007; Yu et al., 2006). To elucidate potential mechanisms underlying the electrophysiological changes observed in CA1 interneurons in our animal model of concussion, we used the NEURON simulation environment (Hines and Carnevale, 1997, 2001), specifically a model of hippocampal PV^+^ basket cells described previously (Bezaire et al., 2016), and adjusted parameters to fit the observed decrease in spike size and firing rate observed in interneurons of mTBI animals (Summary Table 1). The full individual parameter spaces of the CA1 interneuronal model were altered by hand without an *a priori* hypothesis in both the soma and dendrites to examine whether changing multiple parameters might fit the reduced firing rate, spike width and spike amplitude observed in CA1 interneurons post-mTBI (Figure 5). The best fit between model-generated values (Figure 5A, green) and experimental data of injured CA1 interneurons (Figure 5A, red) was achieved by changing the inactivation of voltage-gated sodium channels (h_inf_) in the model (Summary Table 1). The best fit for the PV^+^ interneuron model corresponds to a decrease in firing rate (model = 60% vs. experimental data = 59%), spike width (model = 28% vs. experimental data = 29%), and spike amplitude (model = 27% vs. experimental data = 54%).

**Figure 5.**
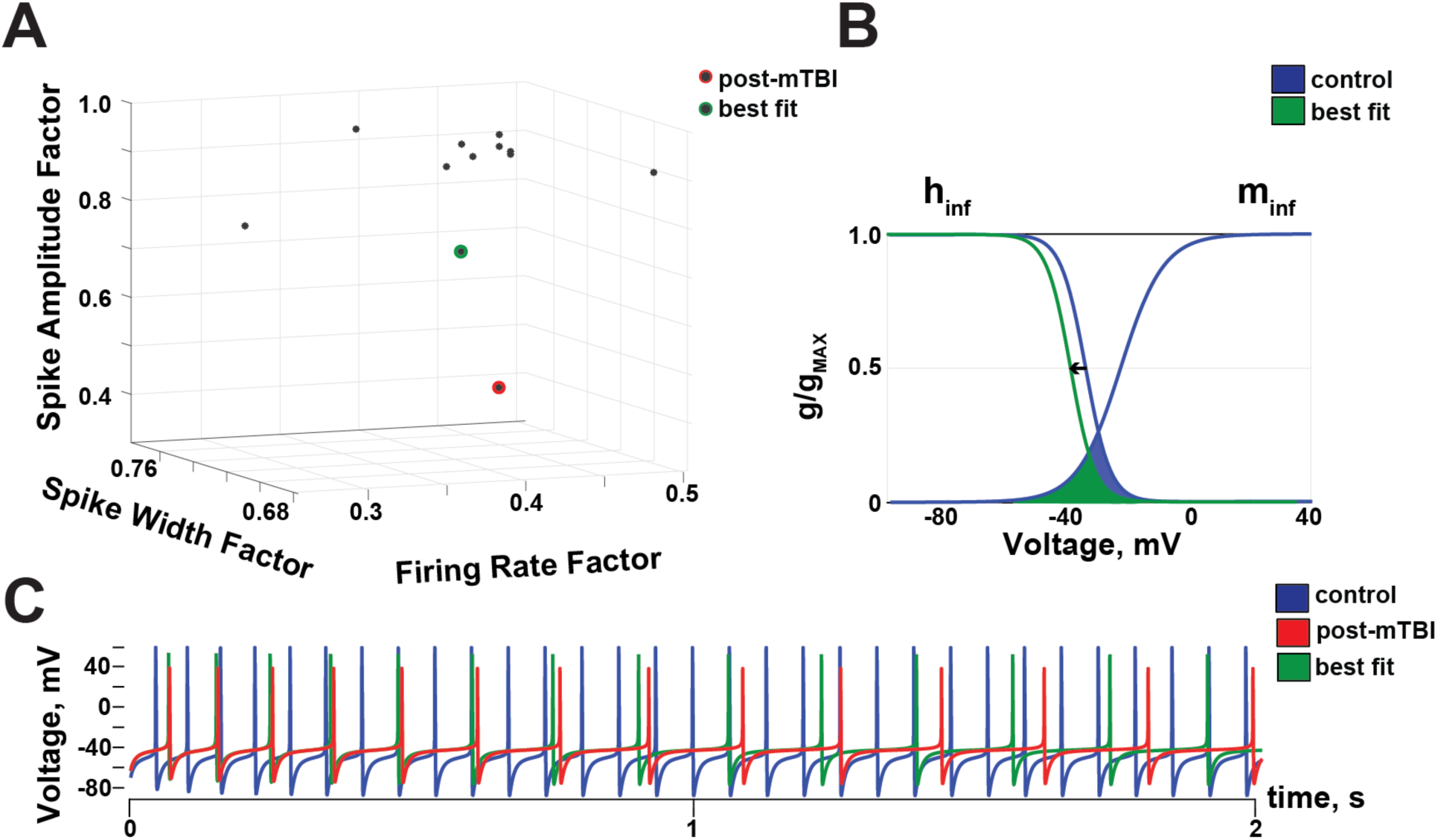
Model of Basket Cells Reveals Potential Dysfunction in Sodium Channels. **A)** NEURON environment was used to model parvalbumin-positive (PV^+^) hippocampal CA1 interneurons (see (Bezaire et al., 2016) for details). The original parameters were altered in both the soma and dendrites to match experimental data from post-mTBI animals (red dot). The best fit (green dot) is achieved by altering a single parameter in the model (th_inf_), and it corresponds to a 60% reduction in the interneuronal firing rate (vs. 59% in the experimental data), a 28% in the spike width (vs. 29% in the experimental data), and a 27% reduction in the spike amplitude (vs. 54% in the experimental data). See Summary Table 1 for details. **B)** Voltage dependence of activation (m_inf_, right curve) and steady-state inactivation (h_inf_, left curve) of sodium currents generated in the model PV^+^ CA1 interneuron (g/g_MAX_ = normalized conductance). A best fit to the experimental data (green) was produced by a negative shift of the inactivation (h_inf_) from –35 mV (blue) to –44.5 mV (green) and is shown with an arrow. Activation (m_inf_) was not altered when using the best fit parameters. The window current of voltage-gated sodium channels (area under the curves) was decreased by ∼ 33% (green). **C)** Changes in firing rate and spike amplitude shown for control (blue), post-mTBI (red), and best fit parameters (green).

**Summary Table 1.**
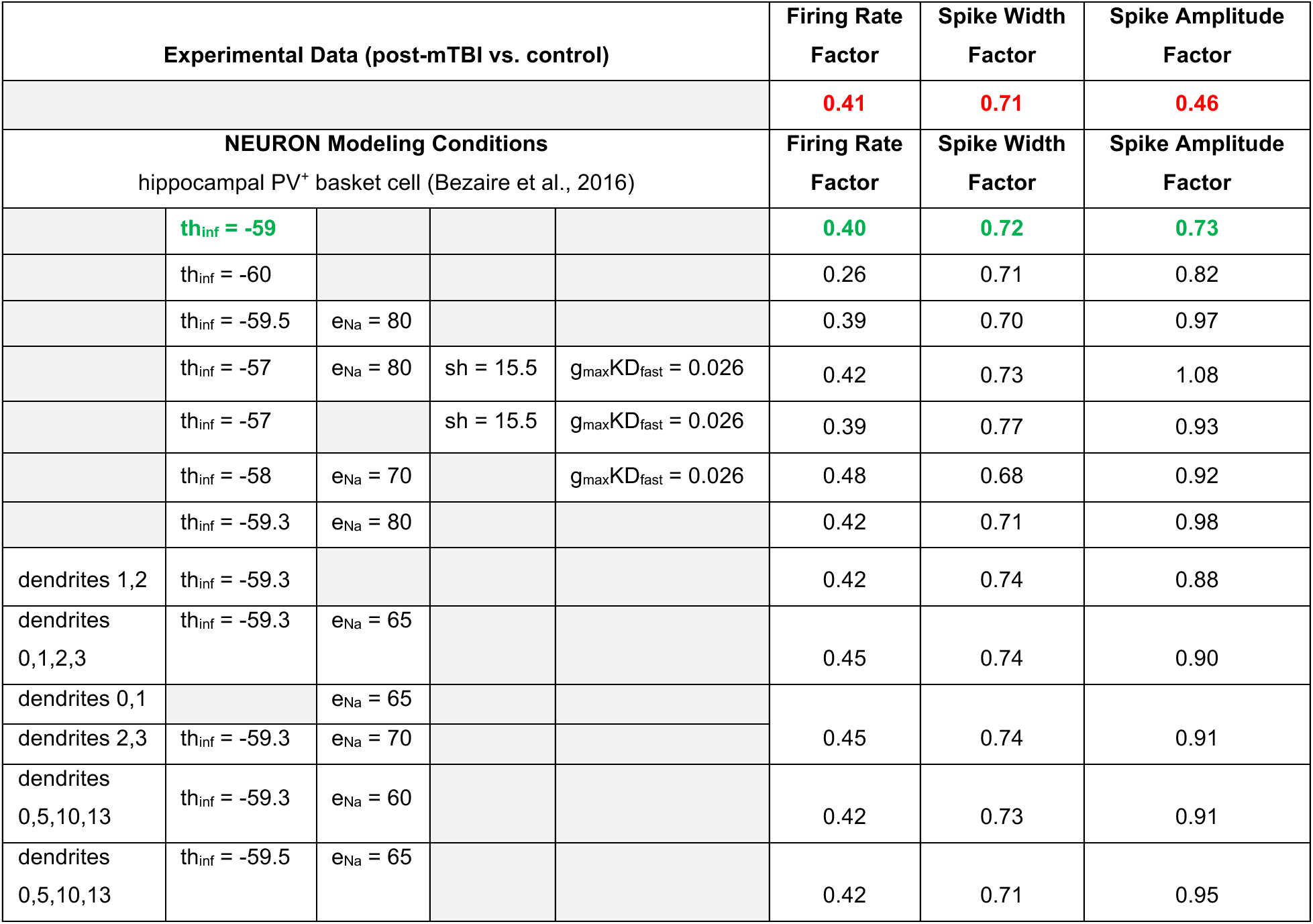
NEURON-simulation environment model of parvalbumin-positive (PV^+^) hippocampal CA1 interneurons. Parameters of the PV^+^ basket cell (Bezaire et al., 2016) were altered to simulate experimental data (red, experimental changes in post-mTBI vs. control groups of animals) observed in CA1 interneurons at 7 days following mTBI. The best fit of the model to the experimental data was achieved by a change of a single parameter, h_inf_ (green).

| Experimental Data (post-mTBI vs. control) |  |  |  |  | Firing Rate Factor | Spike Width Factor | Spike Amplitude Factor |
| --- | --- | --- | --- | --- | --- | --- | --- |
|  |  |  |  |  | 0.41 | 0.71 | 0.46 |
| NEURON Modeling Conditions<br>hippocampal PV<sup>+</sup> basket cell (Bezaire et al., 2016) |  |  |  |  | Firing Rate Factor | Spike Width Factor | Spike Amplitude Factor |
|  | th<sub>inf</sub> = -59 |  |  |  | 0.40 | 0.72 | 0.73 |
|  | th<sub>inf</sub> = -60 |  |  |  | 0.26 | 0.71 | 0.82 |
|  | th<sub>inf</sub> = -59.5 | e<sub>Na</sub> = 80 |  |  | 0.39 | 0.70 | 0.97 |
|  | th<sub>inf</sub> = -57 | e<sub>Na</sub> = 80 | sh = 15.5 | g<sub>max</sub>KD<sub>fast</sub> = 0.026 | 0.42 | 0.73 | 1.08 |
|  | th<sub>inf</sub> = -57 |  | sh = 15.5 | g<sub>max</sub>KD<sub>fast</sub> = 0.026 | 0.39 | 0.77 | 0.93 |
|  | th<sub>inf</sub> = -58 | e<sub>Na</sub> = 70 |  | g<sub>max</sub>KD<sub>fast</sub> = 0.026 | 0.48 | 0.68 | 0.92 |
|  | th<sub>inf</sub> = -59.3 | e<sub>Na</sub> = 80 |  |  | 0.42 | 0.71 | 0.98 |
| dendrites 1,2 | th<sub>inf</sub> = -59.3 |  |  |  | 0.42 | 0.74 | 0.88 |
| dendrites 0,1,2,3 | th<sub>inf</sub> = -59.3 | e<sub>Na</sub> = 65 |  |  | 0.45 | 0.74 | 0.90 |
| dendrites 0,1 |  | e<sub>Na</sub> = 65 |  |  | 0.45 | 0.74 | 0.91 |
| dendrites 2,3 | th<sub>inf</sub> = -59.3 | e<sub>Na</sub> = 70 |  |  |  |  |  |
| dendrites 0,5,10,13 | th<sub>inf</sub> = -59.3 | e<sub>Na</sub> = 60 |  |  | 0.42 | 0.73 | 0.91 |
| dendrites 0,5,10,13 | th<sub>inf</sub> = -59.5 | e<sub>Na</sub> = 65 |  |  | 0.42 | 0.71 | 0.95 |

As detailed in Summary Table 1, the experimental data is best described by a negative shift of h_inf_ towards more hyperpolarized voltages, with the midpoint of the steady state inactivation curve changing from –35 mV to –44.5 mV (Figure 5B, arrow). Under these modeling conditions, the window current of voltage-gated sodium channels, defined as an overlap between the activation (m_inf_) and inactivation (h_inf_) curves (calculated as area under the curves), is decreased by ∼30% following inertial injury. The model output using the best fit parameters is shown in Figure 5C (control – blue, post-mTBI – red, best fit model – green). These results suggest that pathological changes in the inactivation of voltage-gated sodium channels may be a potential mechanism for PV^+^ interneuron dysfunction post-mTBI.

## Discussion

Concussion (or mild TBI) is a prevalent type of brain injury with the potential for long-term consequences (Langer et al., 2020; Veliz et al., 2021). Specifically, concussion is known to lead to acute and chronic cognitive deficits in a subset of mTBI patients, which manifest as poor performance in tests of memory, learning, and delayed recall up to several weeks post injury (Anita et al., 2008; de Freitas Cardoso et al., 2019; Giza et al., 2018). The hippocampus is involved in learning and memory processes (Carr et al., 2011), and its role in integrating information to support these processes may be particularly sensitive to temporal coding disruptions associated with axonal injury or disruptions in oscillatory activity (for review, see (Wolf and Koch, 2016)). Temporal coding in the CA1 region of the hippocampus supports the acquisition, consolidation, and retrieval of spatial and episodic memories by integrating information from entorhinal and CA3 inputs in a temporally precise manner before routing it to downstream structures (Hazan et al., 2006; Koch et al., 2020; Levin et al., 2021; Singh et al., 2014). Changes in neuronal firing and local hippocampal oscillations have been reported in rodent models of mTBI, suggesting abnormalities in temporal coding (Broussard et al., 2020; Ondek et al., 2020; Paterno et al., 2015). However, studies investigating electrophysiological changes following clinical concussion are sparse due to the lack of data from human patients (for review, see (Schmitt and Dichter, 2015)).

Large animal gyrencephalic models offer several advantages in modeling the complex biomechanics and resultant pathophysiology of inertial injury (Smith, 1997), and these models may be the most relevant for understanding the effects of concussion on cognitive processing and memory. We have previously reported changes in hippocampal circuitry following mTBI using an *ex vivo* slice preparation (Wolf et al., 2017). In the current study, we advanced our investigation of electrophysiological changes in CA1 physiology following rotational acceleration injury by performing high-density laminar electrophysiology in the intact anesthetized swine at 7 days post-mTBI. We found that CA1 interneurons had a significantly decreased firing rate, spike amplitude, and spike width, in the injured compared to the control animals. Moreover, we found that concussion resulted in hippocampal hyperexcitability *in vivo*, which confirms findings from prior work utilizing this model (Wolf et al., 2017). Because CA1 activity is known to support learning and memory processes, our combined observations suggest that mTBI-associated changes in CA1 physiology may underlie some aspects of cognitive disruption following mTBI.

### CA1 Interneurons are Preferentially Affected by Concussion

We found that concussion-induced changes in neuronal firing in CA1 were cell-type specific and preferentially affected CA1 interneurons. We showed that concussion, induced by inertial rotation, led to changes in the firing rate and spike shape of CA1 interneurons but not pyramidal cells. Specifically, CA1 interneurons had decreased firing rates at 7 days following concussion, potentially contributing to a decrease in local inhibition resulting in increased excitation and hyperexcitability in the CA1 pyramidal cell layer. Our NEURON modeling results indicated that the decreased firing rate and a reduction in spike amplitude and width observed in CA1 interneurons of mTBI animals may be due to a negative shift in the inactivation of voltage-gated Na^+^ channels. While other groups have proposed a coupled left-shift of activation/availability of sodium channels using an *in vitro* model of stretch injury (Boucher et al., 2012), our computational model did not predict this negative shift in sodium channel activation but instead predicted a negative shift in sodium channel inactivation. Notably, pathological changes affecting the inactivation of voltage-gated Na^+^ channels have been previously reported following injury (Iwata et al., 2004; Wolf et al., 2001). Using an *in vitro* model of axonal stretch injury, we previously demonstrated that traumatic deformation of axons induced abnormal sodium influx through TTX-sensitive Na^+^ channels, leading to an increase in the intracellular Ca^2+^ concentration in axons (Wolf et al., 2001). Furthermore, axonal stretch injury induced structural changes in the alpha-subunit of voltage-gated sodium channels and led to changes in sodium channel inactivation, followed by a long-term increase in the intracellular Ca^2+^ and subsequent proteolysis of the sodium channel alpha subunit (Iwata et al., 2004). Most recently, widespread and progressive loss of nodal axonal sodium channels have been reported in our pig model (Song et al., 2022). These results continue to implicate voltage-gated Na^+^ channel dysfunction as a consequence of TBI, and our translational model of mTBI further suggests that this dysfunction may preferentially affect interneurons in the CA1 region of the hippocampus.

The changes in oscillatory power and entrainment of CA1 neurons in that we observed at 7 days following concussion may also suggest a potential shift in the recruitment/activation of CA1 neuronal subpopulations (Gloveli et al., 2005; Pike et al., 2000). PV^+^ CA1 interneurons (specifically basket cells) are directly involved in feedforward inhibition and have been proposed to stabilize the frequency of local gamma oscillations and maintain a sparsity of neuronal ensembles in the presence of changes in the excitatory drive (Buzsáki and Wang, 2012; de Almeida Licurgo, 2009; Rennó-Costa et al., 2019). This observation may further support TBI associated disruptions in feedforward inhibition provided by PV^+^ interneurons in CA1. This disruption may also impair behavioral tasks requiring network sparsity (e.g., pattern separation), that have been previously reported to be disrupted in a rodent model of mTBI (Munyon et al., 2014). That concussion may lead to interneuronal changes underscores the importance of cognitive tasks that rely on spatiotemporal precision for clinical diagnosis. However, rigorously correlating these changes requires awake behaving electrophysiology and the development of behavioral tasks sensitive to this injury (Ulyanova et al., 2019).

It is worth highlighting that the methods used to distinguish putative CA1 interneurons from putative CA1 pyramidal cells were developed based on activity in the rat hippocampus and are here adapted to the porcine hippocampus for the first time (Broussard et al., 2020; Csicsvari et al., 1999). Electrophysiological characteristics that define spike shape and firing properties of pig hippocampal neurons may vary from those defining rodent CA1 neurons due to the differences in the size of neurons as well as variability in the voltage-gated channels responsible for the generation of action potentials (Sengupta et al., 2013). Also, anesthesia has been known to affect neuronal firing rate as well as the pattern and strength of inputs arriving into the dendritic arbor of the CA1 pyramidal cells (Lustig et al., 2015). These caveats notwithstanding, the relative firing properties and spike shapes of pyramidal cells vs interneurons in CA1 follow the same patterns in swine as in rats with pyramidal cells typically firing in bursts (resulting in a smaller 1st moment of the autocorrelogram (Figure 2A)) and exhibiting lower firing rates and larger spike widths compared to interneurons which have more tonic firing patterns (resulting in a larger 1st moment of the autocorrelogram (Figure 3A)) and exhibit higher firing rates and more narrow spike shapes. Additionally, similarities between the laminar profile of oscillations (theta, gamma and ripple frequency bands) in the porcine and rodent CA1 (Ulyanova et al., 2018) support our use of these sorting techniques in this study despite the novelty of their use in the porcine hippocampus.

### Pyramidal CA1 Cells Are Desynchronized with Respect to Important Organizing Oscillations

Although we found no alterations in intrinsic properties, we demonstrated that CA1 pyramidal cells lose significant synchronization with local gamma oscillations at 7 days following concussion. Previously, we characterized the laminar profile of oscillations in the pig hippocampus (Ulyanova et al., 2018). Here, we observed a significant reduction in gamma oscillation (35 – 75 Hz) power detected locally to CA1 pyramidal cells post-concussion. The reduction in low gamma oscillations (35 – 55 Hz) could be due to a decrease in the fiber volleys of Schaffer collaterals (inputs from the CA3 layer) previously reported in this model (Wolf et al., 2017). Taking into account the fact that networks of interneurons in CA1 support the generation of local high gamma oscillations (55 - 75 Hz) and entrain CA1 pyramidal cells to these oscillations (for review, see (Buzsáki and Wang, 2012; Colgin and Moser, 2010)), these results support our findings above that CA1 interneurons are preferentially affected by inertial injury and that they have not fully recovered by 7 days following concussion. Due to the hypothesized role of theta-gamma coupling in memory and recall, loss of entrainment of CA1 pyramidal cells to gamma oscillations may underlie aspects of post-concussive syndrome (for review, see (Wolf and Koch, 2016)). However, oscillatory activity in this study is influenced by anesthesia, suggesting that these results need to be confirmed in the awake behaving animal.

### Hippocampal CA1 Layer Is Hyperexcitable

In this study, we observed a prolong depolarization in a subset of injured animals (2 out of 9), followed by a loss of synaptic responses to stimulation. In epilepsy, paroxysmal depolarizing shifts and spreading depolarization have been observed in the hippocampus and cortex using animal models as well as in human patients (for review, see (Hotka and Kubista, 2019)). In TBI, spontaneously generated paroxysmal depolarizations have been only detected in organotypic hippocampal cultures following long-term activity deprivation, potentially due to a homeostatic plasticity mechanism (Koch et al., 2010). Therefore, the prolonged depolarizing shift observed using our model of concussion is the first such epileptogenic event described in the *in vivo* preparation. A long-lasting depolarization shift and accompanying repetitive action potentials following paired-pulse stimulation of Schaffer collaterals could be due to changes in fiber volleys, given the differential response to varying stimulation intensities in slice previously reported between control and injured animals using this model (Wolf et al., 2017). Additionally, we also observed recurring paroxysmal oscillatory events in the form of arcuate spikes occurring in a synchronized manner across all hippocampal layers in all post-mTBI but not control animals. Similar epileptogenic spikes have been previously reported after experimental TBI in rodent models and are thought to reflect repeated population spikes associated with hypersynchronous multi-unit discharges (Bragin et al., 2016). Both findings could be substantially influenced by the loss of activity of hippocampal interneurons. Therefore, there are potentially multiple pathological mechanisms involved in the generation of the prolonged depolarizations since we are observing different inputs to the CA1 dendritic arbor simultaneously with these laminar probes.

### Concussion as a Risk for Development of Post-Traumatic Epileptogenesis

Concussion has been previously identified as a risk factor for the development of epilepsy (for review, see (Agrawal et al., 2006)). However, mechanisms underlying the development of post-traumatic epileptogenesis (PTE) post-concussion have not yet been identified. Here, we reported a significant reduction in the power of high frequency oscillations (80 – 300 Hz) detected locally to interneurons (but not pyramidal) CA1 cells, which may be due to the observed reduction in the firing rate of CA1 interneurons or desynchronization of the CA1 network. A reduction in the number of hippocampal interneurons is a prominent pathology associated with the development of unprovoked seizures both in experimental models and in human temporal lobe epilepsy (Huusko et al., 2015). While interneuronal cell loss in our porcine inertial injury model has not been histologically assessed, the number of electrophysiologically active cells recorded in our model was not significantly different between the control and injured animals. Nevertheless, changes in the electrophysiological properties of interneurons may affect circuit level hippocampal interactions and increase the risk for hyperexcitability, and therefore potentially epileptogenesis.

Hippocampal CA1 hyperexcitability and epileptogenesis caused by changes in sodium channels expressed in inhibitory PV^+^ interneurons have been previously reported (Liautard et al., 2013). While histological and electrophysiological studies have previously identified PV^+^ interneurons to be selectively altered in experimental models of rodent TBI (Carron et al., 2020; Figueiredo et al., 2020; MacMullin et al., 2020; Nichols et al., 2018), we could not identify a specific sodium channel responsible for the observed changes in CA1 interneurons following concussion. Using NEURON modeling of a CA1 PV^+^ interneuron (specifically basket cell), we observed a reduction in the sodium channel window current which in turn may also affect resurgent and/or persistent currents generated by voltage-gated sodium channels (for review, see (Eijkelkamp et al., 2012)). Interestingly, changes in the window current of voltage-gated sodium channels have been previously reported in epilepsy (for review, see (Catterall, 2018; Mantegazza et al., 2010)). Potential mechanisms for the negative shift of sodium channel inactivation include altered properties of sodium channel auxiliary subunits (beta (Spampanato et al., 2004) and gamma subunits) as well as the involvement of slow inactivation caused by prolonged depolarization (Vilin and Ruben, 2001), both of which were proposed to play a crucial role in sodium channel associated epilepsies (Catterall, 2018). In addition, multifocal disruption of the blood brain barrier (BBB) has been demonstrated in this model up to at least 72 hours post-mTBI as evidenced by extravasation of serum proteins, fibrinogen, and immunoglobulin G, and could potentially contribute to observed electrophysiological changes (Johnson et al., 2018).

In summary, using a translational, large animal model of concussion we have demonstrated that a single inertial injury alters hippocampal circuitry at 7 days post-injury by affecting the electrophysiological properties of CA1 neurons. In particular, the physiological properties of interneurons in CA1 are substantially altered following concussion. These interneuronal alterations may be due to a change in the inactivation of voltage-gated sodium channels expressed in the somata of PV^+^ basket cells and may underlie the exhibited decrease in local gamma power and the loss of entrainment of pyramidal cells to local gamma oscillations. We also observed hippocampal hyperexcitability in the local field potentials, predominantly following stimulation. These changes may underlie aspects of cognitive dysfunction following concussion, as well as increase the risk for PTE (Lolk et al., 2021). Future studies are needed to investigate if the changes in sodium channel function predicted by our model are observed experimentally. Further assessing these changes over time and determining the differential involvement of specific interneuron populations may reveal therapeutic targets for improving concussion induced memory dysfunction and/or epileptogenesis. Restoring GABAergic interneuronal function before hyperexcitability proceeds to epileptogenesis may improve cognitive outcomes and decrease seizure frequency, as has been previously demonstrated in a mouse model of PTE (Hunt, 2019). While it is unknown whether the observed changes in interneuronal function persist at time points greater than one-week post-injury, pharmacological interventions centered on restoring sodium channel inactivation may be a new target for improving cognitive function following concussion.

## Materials and Methods

### Animal Care

Yucatan miniature pigs (n = 17) were purchased from Sinclair and underwent the current studies at the age of 6 months and a mean weight of 27.7 ± 4.5 kg (mean ± stdev). The animals were fed standard pig chow and water ad libitum. All animals were pair housed when possible and were always in a shared room with other pigs in a research facility certified by the Association for Assessment and Accreditation of Laboratory Animal Care International (AAALAC facility). All experiments were conducted according to the ethical guidelines set by the Institutional Animal Care and Use Committee of the University of Pennsylvania and adhered to the guidelines set forth in the NIH Public Health Service Policy on Humane Care and Use of Laboratory Animals (2015).

### Surgical Procedures

Prior to the procedures, animals were fasted for 16 hours with water remaining ad libitum. After induction with 20 mg/kg of ketamine (Hospira, 0409-2051-05) and 0.5 mg/kg of midazolam (Hospira, 0409–2596-05), anesthesia was provided with using 2-2.5% isoflurane (Piramal, 66794-013-25) via a snout mask and glycopyrrolate was given subcutaneously to curb secretions (0.01 mg/kg; West-Ward Pharmaceutical Corp., 0143-9682-25). The animals were intubated with a size 6.0 mm endotracheal tube and anesthesia was maintained with 2-2.5% isoflurane per 2 liters O_2_. Heart rate, respiratory rate, arterial oxygen saturation, end tidal CO_2_, blood pressure and rectal temperature were continuously monitored, while pain response to pinch was periodically assessed. All these measures were used to maintain an adequate level of anesthesia. A forced air warming system was used to maintain normothermia.

### Closed-Head Rotational Injury using the HYGE Pneumatic Actuator

An injury was induced according to the previously described protocols ((Cullen et al., 2016) for details). Briefly, the injury was performed under isoflurane anesthesia, while physiological parameters of the animals were being closely monitored. Closed-head rotational acceleration was performed using the HYGE pneumatic actuator, a device capable of producing non-impact head rotation (up to 110 degrees in less than 20 msec) with a controlled relationship between maximum rotational acceleration and injury severity. Angular velocity was recorded using a magneto-hydrodynamic sensor (Applied Technology Associates, Albuquerque, NM) connected to a National Instruments DAQ, controlled by LabVIEW. The animal’s head was secured to a padded bite plate under anesthesia and attached to the HYGE device. A single rapid head rotation was performed within a range of rotational velocities (168 – 269 rad/sec, n = 9), with the mean angular peak velocity being 226 ± 13 rad/sec. For a subset of analyses, the injured animals were divided into two subgroups, those injured at ∼190 rad/sec (188 ± 11 rad/sec, n = 4) versus those injured at ∼260 rad/sec (256 ± 7 rad/sec, n = 5). The injury model has been extensively characterized in previously published studies (Grovola et al., 2021; Johnson et al., 2016; Johnson et al., 2018).

#### Control animals

Control animals (total n = 8) included naïve animals (n = 4). Another subset (n = 3) received an additional craniectomy anterior to the recording site as control for another study (ML = 9, AP = +13). A single sham animal underwent all injury procedures 7 days prior to the recording, including attachment to the HYGE device without firing it, and was kept under anesthesia for the same amount of time as injured animals.

### Laminar Multichannel In Vivo Neurophysiological Recordings in Porcine Hippocampus

Electrode implantation was performed in all control and post-mTBI animals (n = 17 total) according to the previously described protocols (Ulyanova et al., 2019; Ulyanova et al., 2018). Briefly, pigs undergoing electrophysiological recordings (control: n = 8 animals; post-TBI: n = 9 animals) were placed in a stereotaxic frame prior to electrode placement. Large animal stereotaxis combined with single-unit electrophysiological mapping via a tungsten monopolar electrode (Cat# UEWSEGSEBNNM, FHC) was used to precisely place a 32-channel laminar silicon probe into the CA1 region of the dorsal hippocampus. For laminar multichannel recordings, either a NN32/LIN linear 32-channel silicon probe (100 μm or 300 μm spacing between individual channels, Cat# P1×32-70mm-100/300-314r-HP32, NeuroNexus) or a NN32/TET custom-design 32-channel silicon probe (275 μm spacing between individual tetrodes, Cat# V1×32-15mm-tet-lin-177, NeuroNexus) was used for electrophysiological recordings (see (Ulyanova et al., 2019) for more details on multichannel silicon probes used in this study). NN32/LIN probes were used in 9 animals (control = 5 vs. post-mTBI = 4), and NN32/TET probes were used in 8 animals (control = 3 vs. post-mTBI = 5). A skull screw was placed over the contralateral cortex as a reference signal.

Laminar oscillatory field potentials and single-unit activity were recorded concurrently in the dorsal hippocampus of pigs under isoflurane anesthesia (2-2.5%). Anesthesia Neurophysiological signals were amplified and acquired continuously at 32 kHz on a 64-channel Digital Lynx 4SX acquisition system with Cheetah recording and acquisition software (Neuralynx, Inc.). The final location of the silicon probes was confirmed electrophysiologically as described previously (Ulyanova et al., 2018). Briefly, the pyramidal CA1 layer was identified by calculating the root mean squared (RMS) power in the 600 - 6,000 Hz frequency band, a proxy for the spiking activity of neurons.

### Electrical Stimulation

In a subset of animals (control = 3, post-mTBI = 7), electrical stimulation was performed in the Schaffer Collaterals using custom-designed concentric bipolar electrodes (Cat# CBBPC75(AU1), FHC) to investigate CA1 responses to the inputs coming in from the hippocampal CA3 layer. A-M Systems Model 3800 stimulator with MultiStim software installed was used to generate electrical pulses, while A-M Systems Model 3820 Stimulus Isolation Unit was used to deliver the current. Input-output stimulation in the range of 100-1000 μA was performed prior to the experimental stimulation for each animal, and the final stimulation amplitude (range 200-500 μA) was selected at the half maximum response to stimulation (also see (Ulyanova et al., 2018) for more details). Stimulation amplitude was not significantly different between the control and post-mTBI groups of animals (control = 325 ± 66 vs. post-mTBI = 386 ± 54 mA, mean ± SEM, p = 0.51).

At the end of electrode insertion procedure, all pigs underwent transcardial perfusion under anesthesia using 0.9% heparinized saline followed by 10% neutral buffered formalin (NBF). Brains were extracted and inspected for gross pathology. Minimal subdural and/or subarachnoid blood regionally associated with electrode insertion was appreciated in all animals. In three animals (control = 1, post-mTBI = 2), minimal subarachnoid blood was also observed ventral to the brainstem. No gross evidence of brain swelling was identified, consistent with prior descriptions of the model.

### The following analyses were performed off-line on the electrophysiological data

#### Spike Detection

Off-line spike detection and sorting was performed on the wideband signals using the Klusta software packages (http://klusta-team.github.io/klustakwik/), and clusters were manually refined with KlustaViewa software (https://github.com/klusta-team/klustaviewa) (Rossant et al., 2016). The number of single units per animal recorded with either type of multichannel silicon probes (NN32/TET vs. NN32/LIN) was not significantly different, with total number of single units recorded in control (n_cells_ = 125) and post-TBI (n_cells_ = 93) groups used for comparison between the groups. The number of recorded cells per animal was not different between the groups (control = 16 ± 3 cells vs. post-mTBI = 10 ± 3 cells, mean ± SEM, p = 0.24). There were no significant differences detected between a number of single units recorded with two types of electrodes (NN32/TET = 12 ± 3 cells vs. NN32/LIN = 14 ± 3 cells per animal, mean ± SEM, p = 0.65) or between a number of single units recorded with each type of electrode for control vs. post-mTBI groups of animals (NN32/TET: control = 16 ± 6 vs. post-mTBI = 9 ± 3 cells per animal, mean ± SEM, p = 0.26; and NN32/LIN: control = 15 ± 5 vs. post-mTBI = 12 ± 6 cells per animal, mean ± SEM, p = 0.69).

Resulting single-unit clusters were then imported into Matlab (version R2023a) for visualization and further analyses using custom and built-in routines (https://www.mathworks.com/products/matlab.html). In order to minimize waveform shape distortion for visualization, the wideband signal was high pass filtered using a wavelet multi-level decomposition and reconstruction filter (level 6, Daubechies 4 wavelet) (Wiltschko et al., 2008). Waveforms were then extracted from this filtered signal and averaged for display. The freely available Matlab packages FMAToolbox (http://fmatoolbox.sourceforge.net (Hazan et al., 2006)) and Chronux (http://chronux.org (Hazan et al., 2006; Mitra, 2007)) were used for analysis of neuronal bursting properties.

### The following electrophysiological characteristics were calculated for each single unit

#### Firing Rate (Hz)

Firing rate was calculated as the average number of spikes per second. All firing rates reported here are of spontaneously active neurons recorded under anesthesia.

#### First Moment of Autocorrelogram (ms)

For each unit, the autocorrelogram was calculated by binning spike times into 1ms bins and computing the correlation of the binned spike times at lags from 1ms to 50ms. The counts in each bin were divided by the total number of counts to create a probability distribution that is used for plotting. The first moment of the autocorrelogram is defined as the mean value of the autocorrelogram in time.

#### Spike Amplitude (μV)

Spike amplitude was calculated as a distance from baseline to the highest peak.

#### Spike Width (ms)

Spike width was calculated at the half point of spike amplitude.

#### Putative Neuron Subtype Classification

Hippocampal CA1 cells were classified into groups of putative pyramidal cells and inhibitory interneurons (for review, see (Wheeler et al., 2015)) following previously described protocols (Broussard et al., 2020; Csicsvari et al., 1999). Briefly, the following criteria were used: 1) <u>anatomical location:</u> only single units from the top portion of the multichannel silicon probe were selected, which anatomically corresponded to the pyramidal CA1 cell layer; 2) <u>firing rate threshold:</u> putative pyramidal CA1 cells had firing rate of 7 Hz or below, while CA1 interneurons had firing rate above 7 Hz (under anesthesia); and 3) <u>spike waveform and autocorrelogram:</u> first moment of autocorrelogram and symmetry of spike waveforms were used to manually place cells into putative CA1 pyramidal cells vs. CA1 interneurons category similarly to previously published criteria for rodent studies (Broussard et al., 2020; Csicsvari et al., 1999).

#### Analysis of Local Field Potentials (LFP)

Acquired wideband LFPs recorded with the 32-channel silicon probe were downsampled to 2 kHz for further analysis. Signals were imported into Matlab software, version R2017a and processed using a combination of custom and modified routines from the freely available Matlab packages FMAToolbox (http://fmatoolbox.sourceforge.net (Hazan et al., 2006)), Chronux (http://chronux.org (Hazan et al., 2006; Mitra, 2007)), EEGLAB (http://sccn.ucsd.edu/eeglab/index.html (Delorme and Makeig, 2004)), and CSDplotter, version 0.1.1 (Pettersen et al., 2006).

#### Entrainment Analysis

Entrainment strength was determined by computing the modulation vector length (MVL) each single unit with oscillations ranging from 1 – 300 Hz on the channel that contained the maximal spike amplitude. Entrainment strength for all single units was then averaged and compared between the control and post-mTBI groups.

#### Power Spectrum Density (PSD)

PSD analysis of hippocampal oscillations was performed for all channels with detectable single units as previously described (Ulyanova et al., 2018). If more than one single unit was recorded on the same channel, the duplicate power spectrum was excluded from the analysis. The calculated PSD was averaged across all animals then z-score normalized for comparative analysis between the control and post-mTBI groups of animals.

### NEURON Modeling

The NEURON simulation environment was used to model the concussion-associated electrophysiological changes observed in CA1 interneurons (Hines and Carnevale, 1997, 2001). Specifically, a model of parvalbumin-positive (PV^+^) basket cells, fast-spiking CA1 interneurons that synapse on the somata and proximal dendrites of CA1 pyramidal cells, was used as described previously (Bezaire et al., 2016; Migliore et al., 1999). The individual parameters of the model (activation / inactivation and the maximum conductance of voltage gated channels, ionic driving force, action potential amplitude, number of dendrites per neuron, etc.) were altered either alone or in a combination with other parameters in a blinded manner to fit the experimental data (summarized in Table 1). The action potentials of the model-generated interneurons were generated and then compared to the experimentally observed values of the firing rate, spike amplitude and spike width following concussion. The parameter(s) most likely to be affected by the injury were identified based on the best fit between experimentally observed data and the NEURON model generated values.

### Statistics and Reproducibility

Electrophysiological data were collected in 17 animals total (control = 8, post-mTBI = 9). Total number of single units in control (n_cells_ = 125) and post-TBI (n_cells_ = 93) groups were used for comparison. The data were analyzed using Matlab (version R2023a, custom and built-in routines) as well as Graphpad Prism (version 10) software. Non-parametric Mann-Whitney test was used for comparative analyses of neuronal properties (firing rate, spike width, and spike amplitude) between the control and post-mTBI groups of animals. Data were displayed as mean ± SEM. For a subset of analyses of neuronal properties, post-mTBI animals were divided into two subgroups, injured at ∼190 rad/sec (n = 4) vs. at ∼260 rad/sec (n = 5). Ordinary one-way ANOVA was used for comparative analysis between the control, ∼190 rad/sec, and ∼260 rad/sec groups of animals. For power and entrainment analyses, two sample t-test was used for comparison, and data were presented as mean ± STD. Threshold for significance was used with p-value of less than 0.05.

